# Conserved tau microtubule-binding repeat histidines confer pH-dependent tau-microtubule association

**DOI:** 10.1101/486688

**Authors:** Rabab A. Charafeddine, Wilian A. Cortopassi, Parnian Lak, Matthew P. Jacobson, Diane L. Barber, Torsten Wittmann

**Affiliations:** Department of Cell & Tissue Biology, University of California, San Francisco 513 Parnassus Avenue; San Francisco, CA 94143; Department of Pharmaceutical Chemistry, University of California, San Francisco 600 16^th^ St; San Francisco, CA, 94158

**Author notes:** To whom correspondence should be addressed, Department of Cell & Tissue Biology, University of California, San Francisco, 513 Parnassus Avenue; San Francisco, CA 94143-0512, USA.

**Keywords:** MAPT, tau, microtubule, neuronal cytoskeleton, intracellular pH, histidine, cancer

## Abstract

Tau, a member of the MAP2/tau family of microtubule-associated proteins, functions to stabilize and organize axonal microtubules in healthy neurons. In contrast, tau dissociates from microtubules and forms neurotoxic extracellular aggregates in neurodegenerative tauopathies. MAP2/tau family proteins are characterized by three to five conserved, intrinsically disordered repeat regions that mediate electrostatic interactions with the microtubule surface. We use molecular dynamics, microtubule-binding experiments and live cell microscopy to show that highly conserved histidine residues near the C terminus of each MT-binding repeat are pH sensors that can modulate tau-MT interaction strength within the physiological intracellular pH range. At lower pH, these histidines are positively charged and form cation-π interactions with phenylalanine residues in a hydrophobic cleft between adjacent tubulin dimers. At higher pH, tau deprotonation decreases microtubule-binding both in vitro and in cells. However, electrostatic and hydrophobic characteristics of histidine are required for tau-MT-binding as substitution with constitutively positively charged, non-aromatic lysine or uncharged alanine greatly reduces or abolishes tau-MT binding. Consistent with these findings, tau-MT binding is reduced in a cancer cell model with increased intracellular pH but is rapidly rescued by decreasing pH to normal levels. Thus, these data add a new dimension to the intracellular regulation of tau activity and could be relevant in normal and pathological conditions.

---

Neuronal development and function relies on a class of structural microtubule (MT)-associated proteins (MAPs) that share conserved MT-binding motifs. These MAP2/tau family proteins associate with MTs in an extended conformation along the MT protofilament ridge (1, 2), and binding is thought to be mediated largely through electrostatic interactions of basic amino acids distributed throughout the MAP2/tau family MT-binding repeats with the negatively charged MT wall. Tau (MAPT), the most intensely studied member of the MAP2/tau family, is expressed in both the developing and adult nervous system, and different tau splice variants contain either three or four MT-binding repeats (3). In adult neurons, tau-MT binding is restricted to the axon. Tau is thought to influence MT stability and mechanics and to regulate axonal transport potentially by organizing inter-MT spacing in the axonal MT bundle (4–6), but its physiological function and regulation is still poorly understood.

Tau is also a key player in progressive neurodegenerative diseases, such as Alzheimer’s disease and frontotemporal dementia. These and other tauopathies are characterized by a loss of axonal tau-MT association and an abnormal extracellular accumulation of aggregated tau (7, 8) although it remains controversial if these tau neurofibrillary tangles are a cause or consequence of neurodegeneration. Both the spatial control of tau-MT binding in normal neurons and the pathological dissociation from MTs highlight the importance of understanding how tau-MT-binding is controlled. Tau has many phosphorylation sites and as is the case with other MT-binding proteins, tau phosphorylation decreases its affinity for MTs, and pathological tau aggregates are formed by all splice variants of hyperphosphorylated tau (8, 9).

However, the intracellular environment is complex and many incompletely understood factors contribute to the control of protein interactions and functions (10). One such underappreciated regulator is intracellular pH (pH_i_) (11). In contrast to previous static views of pH_i_ homeostasis, we now know that pH_i_ is dynamic and changes, for example, during cell cycle progression (12, 13), cell adhesion (14, 15), and migration (16–18). In addition, dysregulated pH_i_ is a feature of many diseases, including cancer (19, 20) and neurodegenerative disorders (21, 22). Although protein pH sensitivity is generally mediated by histidines with a pK_a_ near neutral, other titratable residues can also play critical roles depending on the protein energy landscape and cooperativity (11). Consequently, several examples have emerged in which histidine protonation dynamics regulate protein-protein (23, 24), protein-nucleotide (25), and protein phospholipid interactions (26, 27). Because all MAP2/tau family MT-binding repeats contain invariant histidine residues, we used *in silico*, *in vitro* and *in vivo* methods to ask if tau-MT binding is sensitive to changes in pH_i_. Consistent with the modulation of electrostatic interactions, we find that tau-MT-binding is decreased at increased pH, which opens new directions for understanding tau-MT binding dynamics in normal and pathological cell behaviours.

## RESULTS

### Conserved histidine residues contribute to tau-MT interactions

New cryo-electron microscopy (cryo-EM) for the first time directly resolves the structure of MT-bound tau at near atomic resolution (28). These data indicate that tau-MT interactions extend beyond the central KxGS MT-binding repeat motif. In particular, the structure of the second tau MT-binding repeat (R2) shows that a highly conserved histidine residue, His299, near the R2 C-terminus may contribute to tau-MT binding. His299 points into a hydrophobic cleft defined by β-tubulin Phe399 and Phe395 near the interdimer interface, and protonated His299 forms a T-shaped cation-π interaction with Phe399 (Fig. 1C). Cation-π interactions are common in proteins (29, 30), and histidine-aromatic complexes are energetically favourable in the hydrophobic protein interior (31).

**Figure 1.**
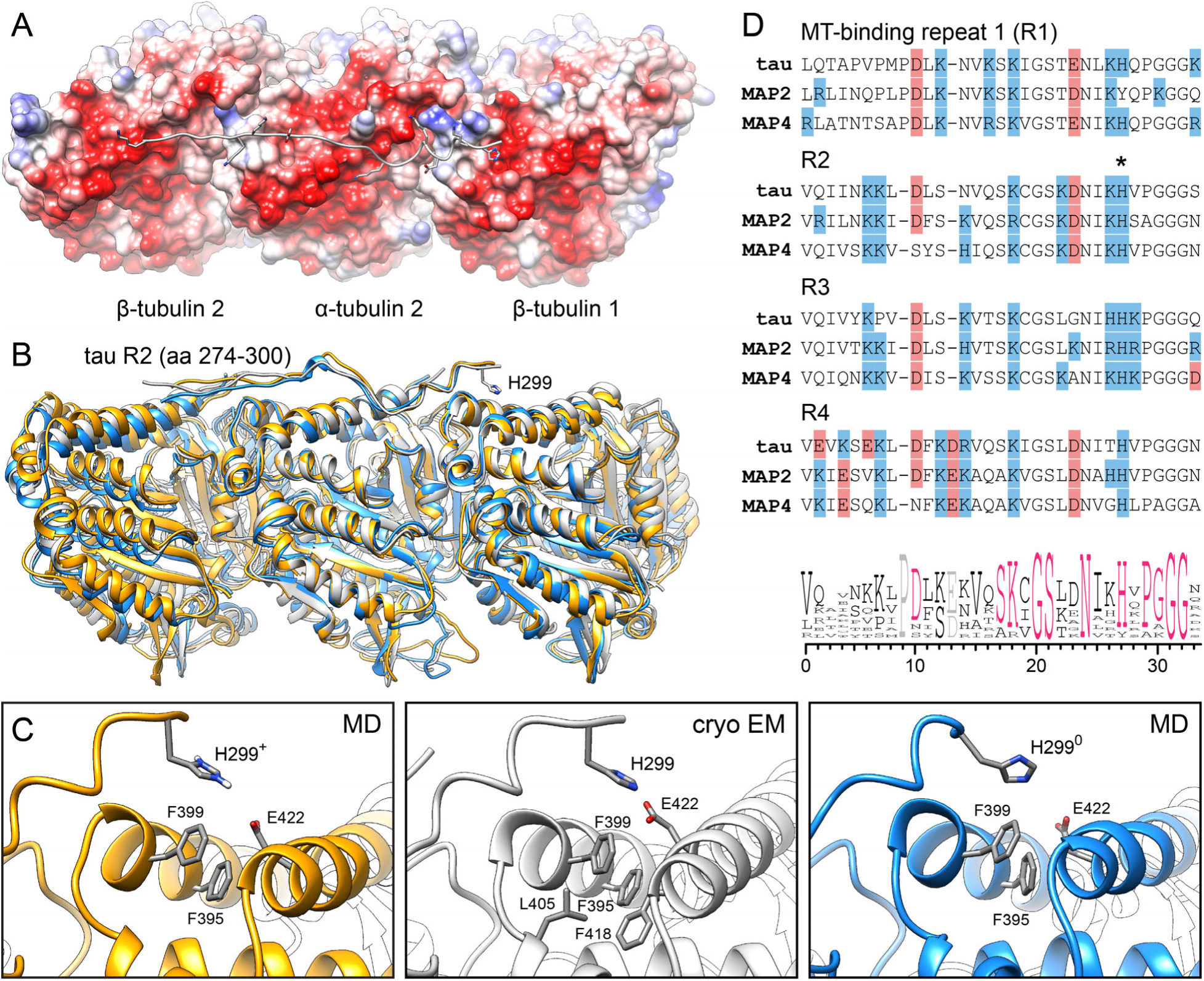
Molecular dynamics simulation of protonation effects on tau His299 interaction with the MT surface. (A) View of the MT-bound tau R2 MT-binding repeat from the MT outside based on the recent cryo-EM structure (PDB ID: 6CVN). The tubulin surface is colored by electrostatic potential highlighting the strongly negative surface charge or the protofilament ridge. (B) Last frames of 5 ns MD simulations of protonated His299^+^ (i.e. low pH; orange) and unprotonated His299^0^ (high pH; blue) compared with the cryo-EM structure (grey). A side view of the tubulin protofilament is shown with tau R2 on top. (C) Close-up views of the binding pocket surrounding His299. (D) Sequence alignment of MT-binding repeats in all members of the human MAP2/tau family indicating the high degree of conservation of the histidine residue in the position equivalent to His299 in tau R2 (asterisk).

Due to the negative electrostatic potential of the MT surface (Fig. 1A), the pK_a_ of His299 is estimated to be upshifted to about 7.0 when compared with the imidazole group in solution that has a pK_a_ of 6.3. A positively charged His299^+^ imidazolium cation is stabilised by favourable electrostatic interactions with negatively charged β-tubulin Glu442 but is also within reach of other hydrophobic contacts (< 7 Å) with β-tubulin residues near the inter-tubulin dimer cleft (Fig. 1C). Based on this analysis, we predicted that His299 is positively charged at neutral pH. His299 protonation could thus be regulated through changes in pH that fall within the physiological range. In contrast, solvent-exposed lysine and arginine residues in the MT-binding repeats have pK_a_ values >10 and are always protonated near neutral pH values. We therefore asked how positively charged His299^+^ or neutral His299^0^ could affect the interaction between tau R2 and MTs by *in silico* molecular dynamics (MD).

Based on the cryo-EM structure, we built models of the tau R2-MT complex with different His299 protonation states. Both systems were then evaluated in short time MD simulations (5 ns). In the model containing positively charged His299^+^, the T-shaped cation-π interaction with Phe399 was maintained throughout the entire simulation (Fig. 1C; supplementary Video S1), with distances not exceeding 5 Å and in agreement with the cryo-EM structure. In contrast, neutral His299^0^ turned away from the hydrophobic cleft and remained in this position during the MD simulation. This also caused higher deviation from the tau-MT cryo-EM structure (RMSD = 2.9 Å) when compared with His299^+^ (RMSD = 1.9 Å) as well as globally increased conformational fluctuations in both tubulin and tau (supplementary Fig. S1). Although our simulations analysed only the one of the 3–4 tau MT-binding repeats for which a cryo-EM structure was available, the MT-binding repeat histidine residues in positions equivalent to His299 are highly conserved (Fig. 1D) (1, 2). In addition, 3R and 4R tau isoforms bind MTs in a similar manner utilizing the full length of the respective MT-binding repeat region (32). Because individually weak MT-binding repeats act together to support high affinity tau-MT interactions (33), histidine deprotonation at increased pH in multiple repeats may thus cooperatively weaken the tau-MT interaction and consequently destabilize the tau-MT complex.

### Increased pH decreases tau-MT binding

To test the hypothesis that increased pH weakens tau-MT binding, we first used an *in vitro* MT co-sedimentation assay. The standard PIPES buffer used in such assays has a pK_a_ of 6.8, which is at the low end of the physiological pH range, hence, we instead incubated 100 nM purified 0N3R tau protein that contains three MT-binding repeats with 0.5 µM paclitaxel-stabilized MTs in a MOPS-based buffer (pKa = 7.2) that provides better buffering capacity in the relevant range. MT-bound tau was then separated from soluble tau protein by centrifugation through a glycerol cushion. Nearly all tau bound to MTs at pH 7.1 (99.5 ± 0.2%; mean ± SD; Fig. 2A). In contrast, the amount of tau in the MT pellet was significantly reduced at pH 7.8 to 91.7 ± 3.4%, and a detectable fraction of unbound tau remained in the supernatant.

**Figure 2.**
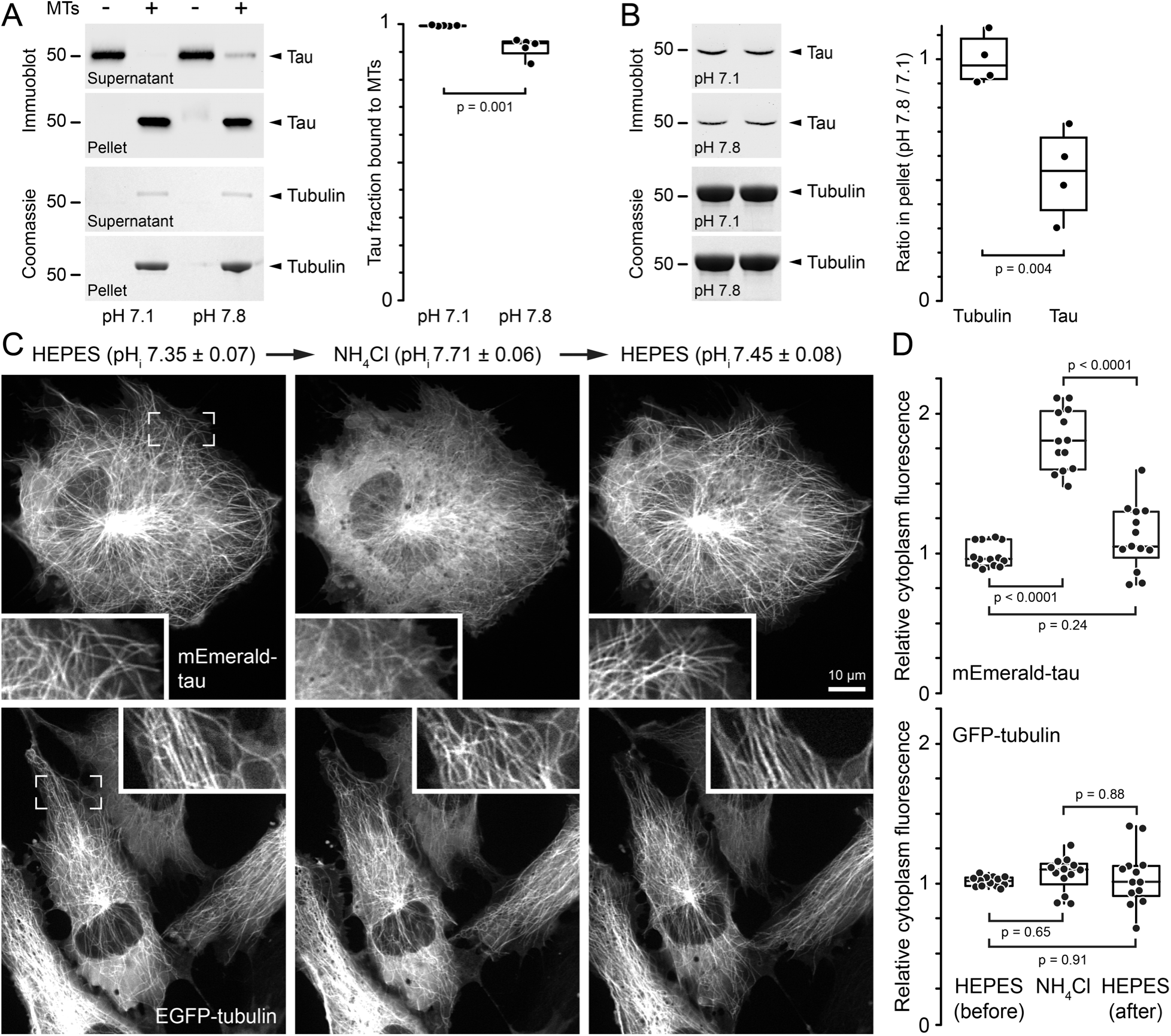
Elevated pH decreases tau binding to MTs. (A) Co-sedimentation assay of 100 nM 0N3R tau protein with 0.5 µM paclitaxel-stabilized MTs. Shown are immunoblots for tau and Coomassie-stained gels for tubulin in the supernatant and pellet at the indicated pH values. Note that tau remains in the supernatant in the absence of MTs at both pH values. The boxplot shows a quantification of the tau fraction recovered in the pellet (*n* = 5). (B) Co-sedimentation assay of 10 nM 0N3R tau protein with 1 µM paclitaxel-stabilized MTs. Because tau is too diluted to be accurately detected in the supernatant only pellets of two independent experiments at the indicated pH values are shown here. The boxplot shows the ratio of either tubulin or tau in the pellet at pH 7.8 compared with pH 7.1 (*n* = 4). Equivalent amounts of supernatant and pellet from different experiments were run in adjacent lanes on the same gels to minimize experimental error. (C) RPE cells transiently expressing either fluorescently tagged tau or tubulin as indicated treated with 20 mM NH_4_Cl to acutely increase pH_i_ in the cytoplasm. Insets show highlighted regions at higher magnification. (D) Quantification of the mEmerald-tau or EGFP-tubulin fluorescence in the cytoplasm (*n* = 13 cells each). Note that elevated pH increases mEmerald-tau signal in the cytoplasm almost 2-fold, which indicates dissociation from MTs. Box plots show median, first and third quartile, with whiskers extending to observations within 1.5 times the interquartile range, and all individual data points. Statistical analysis by Tukey-Kramer HSD test.

Because it is difficult to measure binding differences at near saturation levels, we repeated this experiment with a 20-fold lower tau to MT ratio (Fig. 2B). At these low tau concentrations, it was not possible to accurately determine the amount of tau remaining in the supernatant, but the amount of tau in the MT pellet was reduced to 52.8 ± 18.3% at pH 7.8 compared with pH 7.1. In contrast, the amount of MTs in the pellet remained unchanged (99.5 ± 10.1%) indicating that this difference reflects tau-MT-binding and not a decrease in MT stability at higher pH.

However, *in vitro* experiments cannot fully reconstitute the intracellular milieu or the effect of posttranslational modifications (10). In addition, taxanes used to stabilize MTs *in vitro* alter tau-MT interactions (34). We therefore next asked whether changes in pH_i_ alter tau-MT binding in human retinal pigment epithelial (RPE) cells transiently expressing mEmerald-tagged tau (Em-tau) 0N3R at a low level that did not induce MT bundling. We measured pH_i_ in parallel in untransfected cells loaded with the pH-sensitive dye 2,7-biscarboxyethyl-5(6)-carboxyfluorescein (BCECF) (35, 36). In control pH 7.4 HEPES buffer, the pH_i_ of RPE cells was ~7.4 and Em-tau clearly bound along MTs although a substantial fraction of Em-tau remained in the cytoplasm. Acute elevation of pH_i_ to ~7.7 by addition of 20 mM NH4Cl resulted in a sharp decrease in MT-bound tau (Fig. 2C; supplementary Video S2). Because of low amounts of MT-bound tau at high pH_i_ and movement of the MT network in these time-lapse sequences, direct measurements of MT-bound tau were difficult and unreliable. Therefore, we instead quantified relative Em-tau fluorescence intensity in the cytoplasm, which as expected increased when Em-tau dissociated from MTs in response to experimentally increased pH_i_ (Fig. 2D). Em-tau dissociation from MTs occurred within seconds and was rapidly reversed when the NH4Cl solution was washed out and replaced with control HEPES buffer.

In contrast, the MT network was not affected by these acute pH_i_ changes as quantified by measuring the cytoplasm fluorescence intensity of EGFP-tagged tubulin not incorporated into MTs (Fig. 2C, D). This demonstrates that Em-tau dissociation from MTs at elevated pH_i_ was not a consequence of MT depolymerization, and that the overall polymerization state of the MT network is insensitive to short-term pH_i_ changes within this range.

We observed a similar rapid and reversible disruption of Em-tau binding to MTs when pH_i_ was acutely increased with 100 mM NaCl (Fig. 3), which osmotically activates proton efflux by the plasma membrane Na^+^-H^+^ exchanger NHE1 (35, 36). Tau belongs to the MAP2/tau family of microtubule-associated proteins (MAPs) that share the same repeat motif in the MT-binding domain (1), and the histidine residues at the C-termini of MT-binding repeats in all these proteins are remarkably conserved (Fig. 1D). To test whether other MAP2/tau family proteins respond to intracellular pH_i_ changes, we transiently expressed the MT-binding region of mouse MAP4 containing three MT-binding repeats in RPE cells. Like Em-tau, Em-MAP4 responded to hyperosmotic pH_i_ increase with a reversible decrease in MT-binding (Fig. 3) indicating that decreased MT binding at increased pH_i_ may be a common characteristic of MAP2/tau family proteins.

**Figure 3.**
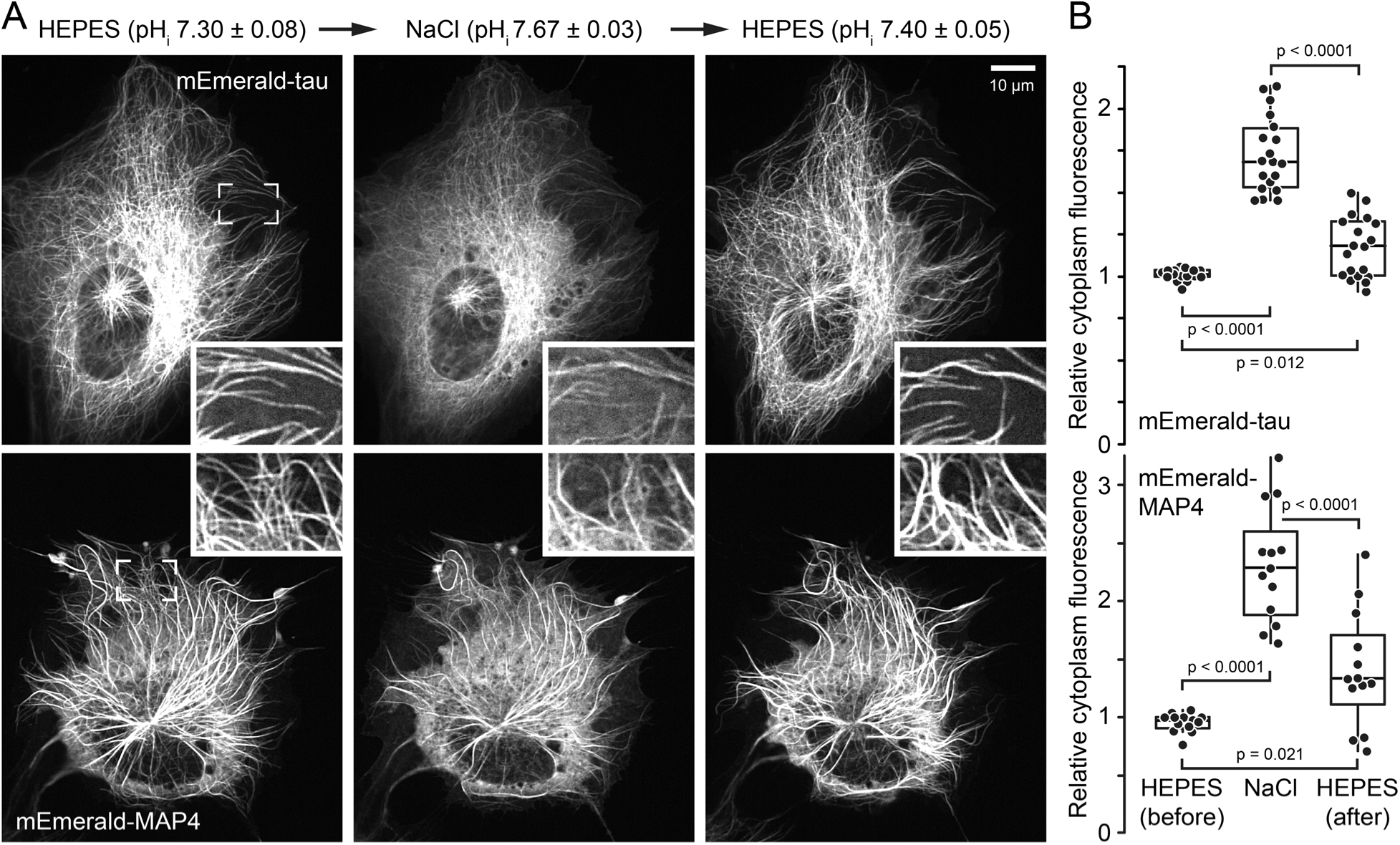
Elevated pH_i_ decreases binding of MAP2/tau family proteins to MTs. (A) RPE cells transiently expressing either mEmerald-tagged tau or MAP4 as indicated treated with 100 mM NaCl to acutely increase pH_i_ in the cytoplasm. Insets show highlighted regions at higher magnification. (B) Quantification of the mEmerald-tau or -MAP4 fluorescence in the cytoplasm (tau: *n* = 19; MAP4: *n* = 13 cells). Both MAP2/tau family proteins reversibly dissociate from MTs at increased pH. Note that MAP4 expression generally results in some degree of MT bundling, and MAP4 does not completely disappear from these bundles at increased pH. Box plots show median, first and third quartile, with whiskers extending to observations within 1.5 times the interquartile range, and all individual data points. Statistical analysis by Tukey-Kramer HSD test.

The above experiments relied on acute changes in pH_i_ that occur within seconds to minutes. To test how long-term pH_i_ changes affected tau-MT interactions, we transiently expressed Em-tau in MCF10A mammary epithelial cells stably expressing an estrogen receptor-induced oncogenic H-RasV12 (ER-RasV12). We previously showed that tamoxifen-induced RasV12 expression increased pH_i_ to almost 7.7 within 24 hours (37). We found that Em-tau was predominantly cytosolic in tamoxifen-induced RasV12-expressing cells. However, acutely decreasing pH_i_ with a pH 7.2 buffer containing the protonophore nigericin, which equilibrates extracellular and intracellular pH, resulted in a rapid rescue of Em-tau binding to MTs (Fig. 4). Together these data demonstrate that an increase in pH_i_ within the physiological range weakens tau-MT binding, which in cells results in substantially less MT-associated tau.

**Figure 4.**
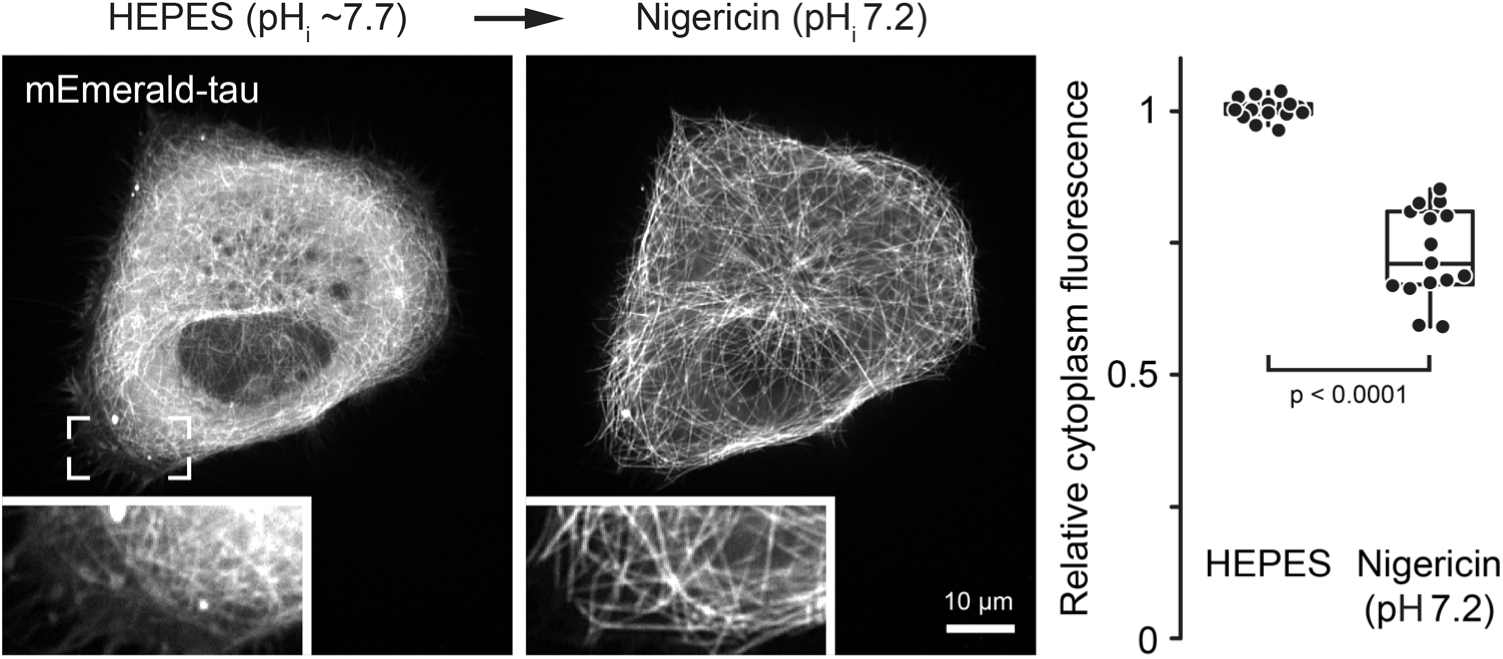
Decreasing pH_i_ in cancer cells rescues tau binding to MTs. (A) MCF10A mammary epithelial cells expressing oncogenic H-RasV12 before and after treatment with a nigericin buffer to equilibrate pH_i_ to 7.2. Inset shows highlighted regions at higher magnification. (B) Quantification of the mEmerald-tau fluorescence in the cytoplasm (*n* = 15 cells). Box plots show median, first and third quartile, with whiskers extending to observations within 1.5 times the interquartile range, and all individual data points. Statistical analysis by Tukey-Kramer HSD test.

### Histidines confer pH sensitivity and are required for interactions with MTs

The location of the His299 cation-π interaction within a hydrophobic pocket indicates that hydrophobic interactions of the histidine aromatic ring are important for MT binding, and may explain the high degree of conservation of these histidine residues. To further investigate this, we sought to identify mutations that mimic His299^0^ and His299^+^ in a pH-independent manner. We therefore built a model in which His299 was changed to a constitutively positively charged lysine residue that we hypothesized could mimic His299^+^. However, MD simulations showed that a lysine substitution behaved more like deprotonated His299^0^ (supplementary Fig. S1). While protonated lysine cation-π interactions are strong and commonly observed in proteins (31), due to the high entropic desolvation cost, the β tubulin hydrophobic cleft near Phe395 and Phe399 cannot readily accommodate the hydrophilic lysine ammonium cation. In fact, Lys299 turns away from the hydrophobic cleft although it can maintain electrostatic interactions with Glu422 (Fig. 5A).

**Figure 5.**
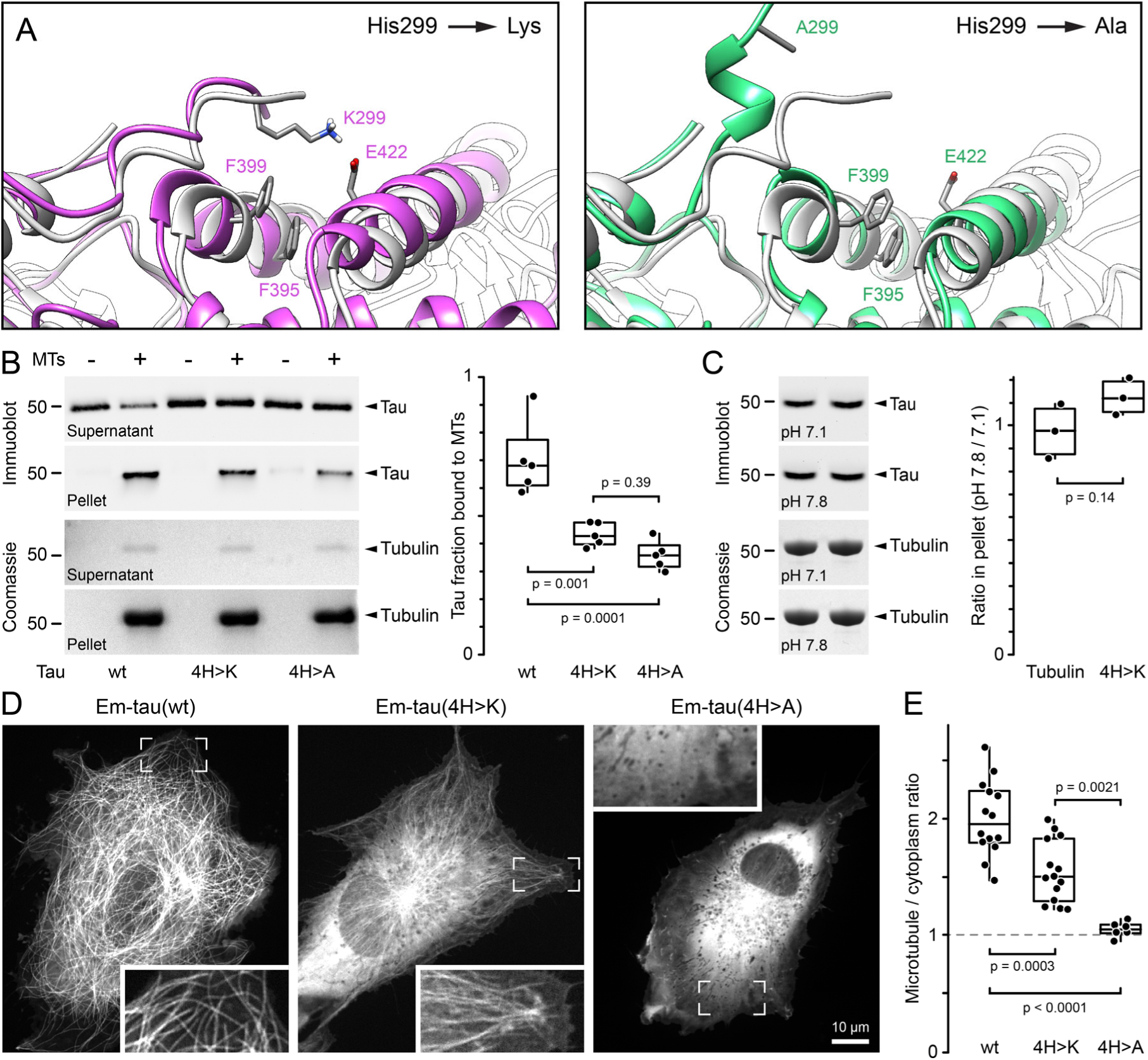
Conserved histidine residues are required for tau binding to MTs. (A) Close-up views of 5 ns MD simulations in which tau R2 His299 was substituted with either lysine or alanine as indicated overlaid with the cryo-EM structure shown in grey. (B) Co-sedimentation assay of 100 nM tau protein with the indicated substitutions of conserved histidine residues with 1 µM paclitaxel-stabilized MTs at pH 6.8. Shown are immunoblots for tau and Coomassie-stained gels for tubulin in the supernatant and pellet. Note that tau remains in the supernatant in the absence of MTs. The boxplot shows a quantification of the tau fraction recovered in the pellet (*n* = 5). (C) Co-sedimentation assay of 10 nM tau(4H>K) protein with 1 µM paclitaxel-stabilized MTs. Pellets from two independent experiments at the indicated pH values are shown. The boxplot shows the ratio of either tubulin or tau(4H>K) in the pellet at pH 7.8 compared with pH 7.1 (*n* = 3). Equivalent amounts of supernatant and pellet from different experiments were run in adjacent lanes on the same gels to minimize experimental error. (D) RPE cells transiently expressing the indicated mEmerald-tagged tau constructs. Insets show highlighted regions at higher magnification. (E) Quantification of the relative enrichment of the indicated mEmerald-tau constructs on MTs compared to cytoplasm signal (wt and 4H>K: *n* = 14; 4H>A: *n* = 6 cells). The dashed line at a MT to cytoplasm ratio of one represents undetectable MT binding. Box plots show median, first and third quartile, with whiskers extending to observations within 1.5 times the interquartile range, and all individual data points. Statistical analysis by Tukey-Kramer HSD test.

To further evaluate the importance of hydrophobic contacts by *in silico* MD, we next analysed a substitution of His299 with alanine, a small non-charged amino acid that is not expected to contribute to either hydrophobic or electrostatic interactions. Like lysine, the alanine substitution also showed increased fluctuations of the tau-MT complex near the C-terminus of tau R2 (supplementary Fig. S1), and throughout the MD simulation Ala299 pointed away from the hydrophobic cleft (Fig. 5A; supplementary Video S3).

While we cannot directly infer binding free energy changes from MD, our *in silico* models support that the positively charged His299^+^ can uniquely form both an electrostatic interaction with Glu422 and a cation-π interaction with Phe399. In contrast, lysine at position 299 only maintains the electrostatic interaction while an Ala299 loses both resulting in partial dissociation of the tau peptide from the MT surface.

To directly test that conserved histidines are required for tau-MT-binding, we generated mutant tau 0N3R protein in which all four histidine residues in the MT-binding repeats (R3 has two adjacent histidines) were substituted with either lysine (4H>K) or alanine (4H>A). If MT-binding relied only on electrostatic interactions with protonated histidines, tau(4H>K) MT-binding would be expected to be similar to wild-type tau, while tau(4H>A) should show reduced MT-binding. Instead, and consistent with our structural analysis, both variants displayed substantially reduced MT binding *in vitro* (Fig. 5B). Although tau(4H>K) seemed to bind to MTs somewhat better than tau(4H>A), *in vitro* this difference was not statistically significant. If these conserved histidines are responsible for the pH sensitivity of tau-MT binding, because of the high pK_a_ of lysine, residual Tau(4H>K) binding to MTs should no longer respond to pH changes within the physiological range. Indeed, in contrast to wild-type tau (Fig. 2B), MT-binding of Tau(4H>K) was not decreased at increased pH in the presence of a 100-fold molar excess of MTs (Fig. 5C). The amount of tau(4H>K) that co-sedimented with MTs at pH 7.8 compared with pH 7.1 was 112.5 ± 8.1%, and not significantly different from the amount of MTs recovered in the pellet at the two different pH values.

Lastly, we tested binding of fluorescently tagged tau variants to MTs in RPE cells expressing equivalent levels of wild-type or mutated Em-tau (Fig. 5D, E; supplementary Fig. S2). In cells, binding of Em-tau(4H>K) to MTs was significantly reduced compared with wild-type tau, and Em-tau(4H>A) did not bind MTs at all. Together, these data confirm that conserved histidine residues in the tau MT-binding repeats are required for tau MT-binding that are not solely explained by electrostatics, and protonation of these histidines can be titrated to modulate tau-MT interactions.

## DISCUSSION

We used multiple approaches including MD simulations and MT binding assays *in vitro* and in cells to show that interactions of MAP2/tau family proteins with MTs are sensitive to pH changes within the physiological intracellular range. We also demonstrate that histidine residues near the C-terminal end of MAP2/tau family MT-binding repeats are essential. Tau mutants with these histidines substituted with either alanine or lysine show reduced or completely absent MT-binding in cells. In addition, at least *in vitro* these conserved histidines mediate pH sensitivity as replacing them with lysine residues, which are insensitive to pH changes in the physiological range, abolishes pH-modulation of MT-binding *in vitro*. However, we were not able to test this rigorously in cells because of the already substantially reduced MT-binding of the H>K mutant.

Although these MAP2/tau MT-binding repeat histidine residues are highly conserved, their importance has not been recognized because the intrinsically disordered nature of MAP2/tau family proteins complicates structural analysis. Previous studies did not resolve MT-bound peripheral regions of these MT-binding repeats and predominantly found MAP2/tau family densities along the protofilament ridge (2). Although a high-definition model of native MT-bound tau is still not available, recent cryo-EM data of MT-bound constructs of human tau MT-binding repeat R2 show a close interaction of His299 with the tubulin interdimer interface (28). At low pH, our structural and MD analysis defines this interaction as a stable cation-π interaction of the protonated histidine with specific aromatic residues in β-tubulin H11. This requires the amphiphilic character of protonated histidine as deprotonation at high pH or substitution with either non-aromatic or non-charged amino acids substantially weakens tau-MT interactions. Thus, our data are consistent with older biochemical crosslinking data showing tau interaction with H11, H12 and the flexible loop connecting these helices (38) and new ultrastructural analysis of MT-bound MAP4 that indicates a strong contribution of the tubulin interdimer interface to the overall MAP4-MT binding energy (39).

Because of its role in neurodegenerative disease, tau is the most widely studied member of the MAP2/tau family, and much research has focused on the role of tau aggregation in Alzheimer’s disease and related tauopathies with over 14000 publications listed in PubMed (3). Less is known about how MAP2/tau family proteins control the MT cytoskeleton in normal neurons. *In vitro*, tau promotes MT polymerization and protects MTs from disassembly likely by spanning multiple tubulin dimers and thus inhibiting their dissociation. Consequently, MAP2/tau family proteins are thought to stabilize MTs in neurons. Recent data of shRNA-mediated tau depletion in primary neurons indicate that tau promotes growth cone MT elongation (40). In addition, tau inhibits EB1 association with growing MT ends (41) suggesting a multiple roles of tau in controlling neuronal MT polymerization dynamics. High levels of MAP2/tau family proteins induce MT bundles in non-neuronal cells with inter-MT spacing similar to neurite MT bundles (6). Binding of MAP2/tau family proteins along MTs also inhibits MT motor movements (5). However, these functional data rely mostly on *in vitro* experiments or MAP2/tau protein overexpression, and to what extent these reflect physiological neuronal tau functions remains unclear. For example, MT-based transport appears unaffected in tau knock-out mice (42), and moderate tau expression levels have little effect on MT network organization. In summary, much remains to be learned about how MAP2/tau proteins control physiologically normal neuronal MT cytoskeleton function. What we do know though is that neurons are morphologically complex and thus the activity of neuronal cytoskeleton proteins must be controlled precisely.

Characteristic of many intrinsically disordered proteins are their complex phosphorylation patterns. MAP2/tau proteins are no exception (3, 43) and phosphorylation within the MT-binding repeats generally reduces tau MT-binding, consistent with electrostatic repulsion between negatively charged phosphates and the strong negative electrostatic field of the MT surface. We propose that the upshifted pK_a_ of the conserved MT-binding repeat histidine residues allows them to act as pH sensors and modulate tau-MT electrostatic interactions within the physiological pH range. Consistent with a decreased tau-MT interaction strength, individual tau molecules also show increased diffusion along MTs at increased pH (44). Thus, pH_i_ changes may add another layer of control to MAP2/tau MT-binding. Observed differences in the magnitude of pH-modulated tau MT-binding between our experiments *in vitro* and in cells may be due to partial phosphorylation of tau in cells. However, tau-binding to MTs is also sensitive to subtle changes in MT lattice geometry (34). Although we find that the total MT polymer is insensitive to short-term pH_i_ changes, we cannot rule out that pH-mediated conformational changes of the MT wall contribute to pH-sensitive tau-MT interactions.

Currently, we can only speculate as to the physiological relevance of pH modulating tau-MT-binding. Because neurons are morphologically differentiated with highly compartmentalized cytoplasm, it is possible that developmental or spatial control of pH_i_ in developing neurons contributes to spatiotemporal control of MAP2/tau activities, but potential local differences in neuronal cytoplasmic pH_i_ remain poorly characterized (45, 46). However, acidification of the neuronal cytoplasm has been associated with and speculated to be a causative agent in neurodegeneration (21). Our current data would predict enhanced tau-MT binding in acidified neurons, which is inconsistent with tau aggregation due to weakened tau-MT interactions in Alzheimer’s disease. Alternatively, increased binding of protonated tau to negatively charged phospholipid membranes may increase toxic effects of aggregated tau (47).

Lastly, ectopic tau expression has been described in many types of cancer and elevated tau levels appear to correlate with aggressive metastatic cancer phenotypes and resistance to taxane chemotherapy (48–50). Cancer cells generally have elevated pH_i_ (19, 20) although a large variability in different types of cancers is likely. Our data indicate that tau-MT-binding is greatly reduced in a breast cancer model driven by oncogenic RasV12 and is rapidly rescued by returning pH_i_ to a physiological normal level. Thus, we would predict that taxane sensitivity positively correlates with pH_i_ in tau-expressing cancers, which could serve as a diagnostic tool for taxane chemotherapy. In conclusion, even though we do not yet understand the functional relevance of pH-modulated tau-MT interactions in normal or pathological conditions, the current study adds another facet to the already complex regulation of MAP2/tau cell biology.

## EXPERIMENTAL PROCEDURES

*Molecular dynamics –* The recently reported cryo-EM structure of MT-bound tau MT-binding repeat 2 (amino acids 274–300; PDB ID: 6CVN) was used as starting point for MD simulations (28). The pK_a_ of MT-bound His299 was predicted with propKa 3.0 software. Two different protonation states for His299 were modeled, a neutral His299^0^ protonated on the ε nitrogen, as predicted by the Maestro Protein Preparation package, and a double protonated form, His299^+^. His299 was also mutated to lysine or alanine using the ‘swappa’ command as implemented in Chimera (51). Preceding MD simulations, the Tau-MT model structures were submitted to 1,000 steps of steepest descent minimization, followed by 1,000 ps of equilibration. The SHAKE algorithm was used to constrain all bonds containing hydrogen atoms. A cut-off of 12 Å was used for long-range interactions. These energy-minimized models were then submitted to short time MD simulations (5 ns) using the NPT ensemble with the Amberff12SB force field (52). The Berendsen pressure coupling scheme was used for keeping the pressure constant at 1 atm. The temperature of the production was constant at 310K using the Langevin thermostat.

*Plasmids and protein production* – Mammalian expression plasmids mEmerald-MAPTau-C-10 and mEmerald-MAP4-C-10 (Addgene Plasmid # 54152) were from the Michael Davidson plasmid collection. Tau(4H>K) and (4H>A) mutations were constructed as gene blocks (Integrated DNA Technologies) and inserted into the wildtype pmEmerald-MAPTau-C-10 plasmid using Gibson assembly.

For bacterial expression, tau constructs were subcloned into pET28a bacterial expression vectors using Gibson assembly containing an N-terminal 6xHis tag. pET28a constructs were expressed in BL21(DE3) *E. coli* cells in Luria Broth after induction with 1 mM isopropyl β-D-1-thiogalactopyranoside (IPTG) overnight at 18°C. Bacteria pellets were lysed by sonication on ice in 50 mM Tris-HCl pH 7.5, 150 mM NaCl, 1 mM DTT and protease inhibitors. Recombinant tau was purified by affinity chromatography on Ni-NTA resin, eluted into 50 mM Tris-HCl pH 7.8, 200 mM NaCl, 200 mM imidazole, then dialyzed into 20 mM Tris-HCl pH 7.4, 150 mM NaCl, 10% glycerol, 1 mM DTT, concentrated to ~50 µM, and flash frozen and stored in aliquots in liquid N_2_.

*In vitro MT-binding assays* – Paclitaxel-stabilized MTs were prepared by polymerizing 25μM purified porcine or bovine tubulin in 80 mM K-PIPES pH 6.8, 5 mM MgCl_2_, 1 mM EGTA, 33% glycerol and 1 mM GTP at 37 °C. After 30 min 50 µM paclitaxel was added. To test MT-binding of histidine mutants, 100 nM tau protein was incubated with 1 µM MTs in 250 µl BRB80 (80 mM K-PIPES pH 6.8, 1 mM MgCl2, 1 mM EGTA) with 1 mM DTT, 10 µM paclitaxel, and 10 µg/ml BSA. For testing pH-sensitive MT-binding *in vitro*, we instead used a MOPS-based buffer with improved buffering capacity in the physiological pH range (20 mM MOPS, 0.1 mM EDTA, 0.1 mM EGTA, 5 mM magnesium acetate, 50 mM potassium acetate) with 1mM DTT, 10 μM taxol, and 10 μg/ml BSA. The MOPS buffer pH was adjusted to 7.2 and 8.0, respectively. pH values reported on the figures were measured after adding buffer equivalent to the amounts of MTs and tau added, which lowered the pH in the binding reactions by 0.1–0.2 pH units. For reactions with 10 nM tau, the volume was doubled to 500 µl.

After 20 min incubation at room temperature, MT-binding reactions were overlaid onto 400–600 µl of BRB80 or the MOPS-based buffer with 10 µM paclitaxel containing 60% w/v glycerol and centrifuged in a Beckman TLA 100.2 rotor at 50,000 rpm for 20 min at 25°C. After centrifugation a portion of the supernatant (S) was saved for analysis, the cushion carefully washed with ddH2O to minimize contamination of the pellet and removed, and the pellet (P) resuspended in SDS-PAGE sample buffer. Equivalent amounts of supernatant and pellet were analyzed by SDS-PAGE on 4–12% NuPage gels (Invitrogen) and immunoblotting with a tau antibody (E-4, sc-515539; Santa Cruz Biotechnology). Coomassie-stained gels and chemiluminescent immunoblots were imaged with a FluorChem Q gel documentation system (92-14116-00, Alpha Innotech). The MT-bound tau fraction was calculated as (I_P_−I_bkg_)/(I_S_+I_P_−I_bkg_).

*Cell culture, live cell imaging and analysis –* Retinal pigment epithelial (RPE) cells were maintained in RPMI medium (Invitrogen) supplemented with 10% fetal bovine serum (Atlanta Biosciences) and 1% Glutamax (Invitrogen) at 37°C, 5% CO2. MCF10A cells stably expressing ER-RASV12 were maintained as described (37). For transient protein expression and microscopy, cells were plated in 35 mm glass-bottom dishes (MatTek), transfected after 24 h using Fugene6 (Promega) and 1 µg plasmid DNA, and used for experiments 24 h later after replacing the transfection medium.

To determine the pH response of tau MT-binding in cells, the tissue culture medium was replaced with 25 mM Na-HEPES pH 7.4, 140 mM NaCl, 5 mM KCl, 10 mM glucose, 1 mM MgSO4, 1 mM K_2_HPO_4_/KH_2_PO_4_ (pH 7.4), and 2 mM CaCl_2_. Cells were imaged on an environmentally controlled spinning disk confocal system as described (53, 54). After 3-5 images in the control HEPES pH 7.4 buffer, the buffer was replaced with the same buffer either containing 20 mM NH_4_Cl or 100 mM NaCl. After 5–10 min the buffer was once again replaced with control HEPES buffer. Intracellular pH changes with these treatments were determined in parallel in untransfected cells loaded with 2,7-biscarboxyethyl-5(6)-carboxyfluorescein (BCECF) essentially as described (35, 36). BCECF fluorescence at λ_Em_ = 530 nm at two different excitation wavelengths (λ_Ex_ = 440 nm and 490 nm) were acquired using a SpectraMax M5 plate reader (Molecular Dynamics, Sunnyvale, CA), and calibrated to cells treated with 10 µM of the proton ionophore nigericin (Invitrogen) at pH 7.5 and pH 6.6.

To indirectly analyze the amount of MT-bound tau at different pH_i_ values, we measured the mEmerald-tau fluorescence in the cytoplasm, which increases if less tau is MT-bound. mEmerald fluorescence intensity was measured in three small regions-of-interest (ROIs) per cell in three different time-points on the unprocessed 16-bit images per buffer condition, and as necessary, the ROIs were moved slightly between images to avoid MT movements. ROI intensities were corrected for photobleaching by normalizing to the total cell fluorescence, and then normalized to the first time point to preserve variability between measurements. The relative amount of MT-bound tau mutants was quantified as described (34). All image analysis was done in NIS Elements version 4.3 or higher (Nikon). Significance of multiple comparisons was calculated by Tukey–Kramer honest significant difference (HSD) test in Analyse-It for Microsoft Excel. Figures were assembled in Adobe Illustrator CS5, and videos were made using Apple QuickTime Pro. Box-and-whisker plots show median, first and third quartile, observations within 1.5 times the interquartile range, and all individual data points.

## Supporting information

## Acknowledgements

We thank Rebecca Heald’s lab for a kind gift of purified tubulin when we ran low, Katherine White for help with the production of recombinant tau protein, and XSEDE SDSC (Comet) for supercomputer facilities.

## Author contributions

R.C., D.L.B. and T.W designed experiments and analyzed data. R.C. performed in vitro biochemistry. D.L.B. performed experiments in cells. P.L. contributed to experiments. W.A.C. and M.P.J. performed molecular dynamics simulations and structural analysis. All authors contributed to writing of the manuscript.

## FOOTNOTES

This research was supported by a Paul G. Allen Frontiers grant to D.L.B., M.P.J. and T.W., by National Institute of General Medical Sciences R01 GM116384 to D.L.B., and by National Institute of Neurological Disorders and Stroke R01 NS107480 to T.W.

The abbreviations used are: MT, microtubule; MAP, microtubule-associated protein; pH_i_, intracellular pH; MD, molecular dynamics

