## Supplementary material for "Conserved tau microtubule-binding repeat histidines confer pH-dependent tau-microtubule association"

### SUPPORTING INFORMATION

**SUPPLEMENTARY VIDEOS:**

**Video S1. Molecular dynamics simulation of the interaction of protonated tau R2 His299$^+$ with the MT surface.** The C-terminus of tau MT-binding repeat R2 remains in close contact with the MT throughout the molecular dynamics experiment. Tau His299$^+$ and β-tubulin residues Phe399 and Glu442 are shown in addition to the backbone ribbons. This video corresponds to Fig. 1C.

**Video S2. Em-Tau MT binding in response to increased intracellular pH.** Time-lapse recording of an RPE cell expressing Em-Tau first in HEPES buffer, then treated with 20 mM $NH_4Cl$, and finally after recovery in HEPES buffer as indicated in the upper right corner. This video corresponds to Fig. 2C. Scale bar, 10 µm. Elapsed time is shown in min:sec.

**Video S3. Molecular dynamics simulation of the interaction of tau R2 in which His299 was replaced with an alanine with the MT surface.** The C-terminus of the mutated tau MT-binding repeat R2 is turned away from the MT surface throughout the molecular dynamics experiment. Mutated tau Ala299 and β-tubulin residues Phe399 and Glu442 are shown in addition to the backbone ribbons. This video corresponds to Fig. 5A.

**SUPPLEMENTARY FIGURES:**

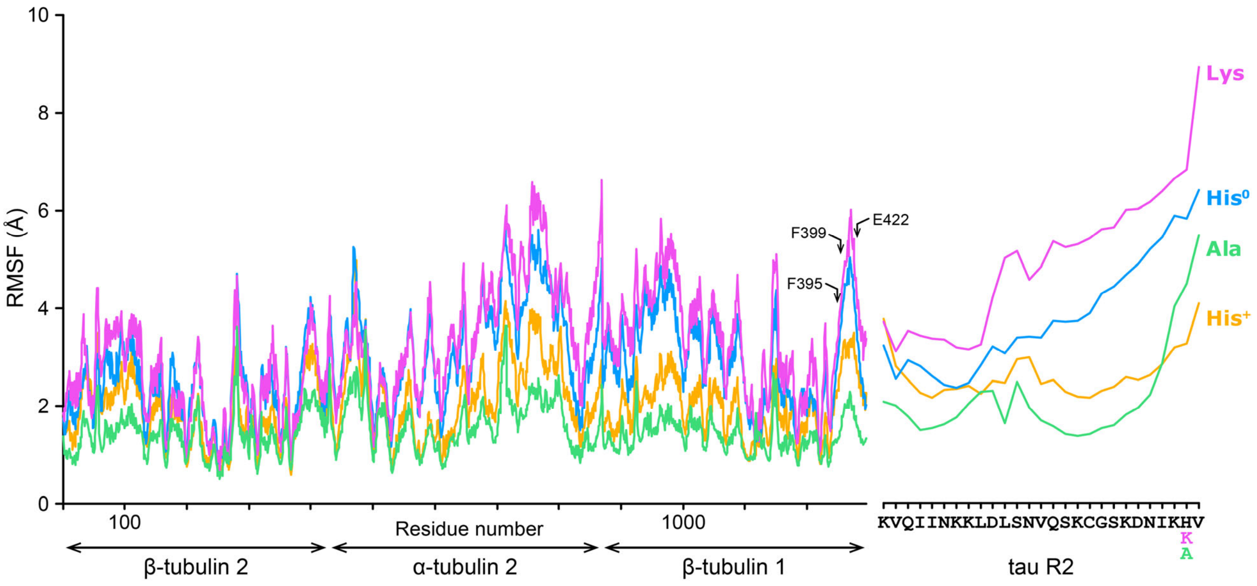


**Figure S1. Conformational fluctuations of the tau-MT complex molecular dynamics simulations.** Graphs show the root mean square fluctuation (RMSF) of the indicated molecular dynamics simulations on a residue-by-residue basis. The MD simulation with protonated $His299^{+}$ shows the lowest RMSF near the tau R2 C-terminus. In contrast, unprotonated His299 or substitution of His299 with either lysine or alanine results in a large RMSF increase at the tau R2 C-terminus. This conformational flexibility also propagates to residues within 5 Å of His299 and the loop between β-tubulin helices H11 and H12 except for the alanine substitution that likely has lost all interaction with the MT surface. Note that the tubulin (left) and tau R2 (right) residue axis is not shown at the same sequence scale.

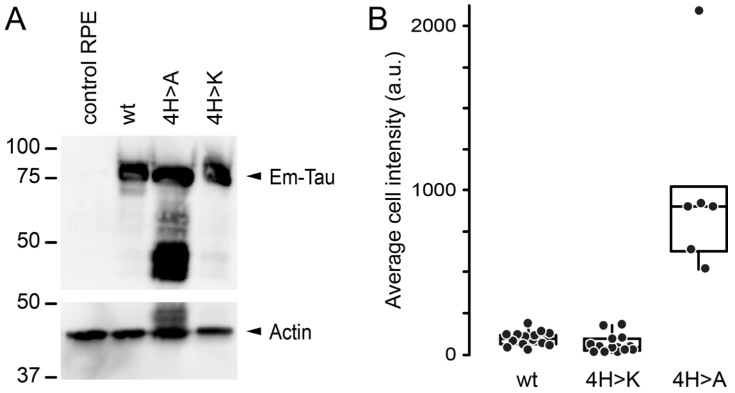


**Figure S2. Transient expression of mEmerald-tagged tau mutants in RPE cells.** (A) Immunoblot of RPE cell lysates transfected with the indicated Em-tau constructs probed with either anti-tau or anti-actin antibodies (clone C4; Millipore MAB1501). Note that all constructs express full-length Em-tau at the expected molecular weight although the 4H>A mutant shows a considerable amount of degradation. (B) Total Em-tau fluorescence intensity in the cells analysed in Fig. 5E indicating higher expression level of the 4H>A mutant.
